# A practical solution for preserving single cells for RNA sequencing

**DOI:** 10.1101/160804

**Authors:** Moustafa Attar, Eshita Sharma, Shuqiang Li, Claire Bryer, Laura Cubitt, John Broxholme, Helen Lockstone, James Kinchen, Alison Simmons, Paolo Piazza, David Buck, Kenneth J. Livak, Rory Bowden

## Abstract

The design and implementation of single-cell experiments is often limited by their requirement for fresh starting material. We have adapted a method for histological tissue fixation using dithio-bis(succinimidyl propionate) (DSP), or Lomant’s Reagent, to stabilise cell samples for single-cell transcriptomic applications. DSP is a reversible cross-linker of free amine groups that has previously been shown to preserve tissue integrity for histology while maintaining RNA integrity and yield in bulk RNA extractions. Although RNA-seq data from DSP-fixed single cells appears to be prone to characteristic artefacts, such as slightly reduced yield of cDNA and a detectable 3’ bias in comparison with fresh cells, cell preservation using DSP does not appear to substantially reduce RNA complexity at the gene level. In addition, there is evidence that instantaneous fixation of cells can reduce inter-cell technical variability. The ability of DSP-fixed cells to retain commonly used dyes, such as propidium iodide, enables the tracking of experimental sub-populations and the recording of cell viability at the point of fixation. Preserving cells using DSP will remove several barriers in the staging of single-cell experiments, including the transport of samples and the scheduling of shared equipment for downstream single-cell isolation and processing.

## Introduction

Single-cell techniques are revolutionising biology by improving the resolution of experiments from the tissue to the cellular level. In particular, the combination of rapid advances in cell isolation technologies, straightforward single-cell RNA sequencing (scRNA-seq) methods, and cheaper high-throughput sequencing, are enabling the capture of detailed information from an ever larger number of individual cells. The Fluidigm C1^™^ nanofluidics system is a widely used platform for the isolation and processing of single cells for an increasing range of genomics applications. The C1 platform benefits from ease-of-use, the ability to image cells after capture, and favourable inter-cell consistency, which reportedly is due to its small reaction volumes (1). A significant unsolved technical issue in single-cell isolation on the C1 (and other platforms) is the maintenance of cell and analyte integrity during preparation of the sample for single-cell isolation. This process can take up to several hours during which the cells are removed from their normal environment. After isolation, molecules such as RNA are typically stabilised by cell lysis in appropriate conditions, so the vulnerable period is the time from initial sample collection to cell lysis. This is a problem that is not unique to single cell studies. Many *in vitro* biochemical analyses suffer from the unresolved concern that manipulation of the sample may be altering the so-called “natural” state of the cells (2). The ability to “freeze” cell processes as early as possible in the experimental protocol will increase researchers’ confidence that observations represent biological rather than technical effects.

Another factor limiting the feasibility of single-cell studies may be availability of the equipment needed at the time when the cells become available for isolation. This can be because of scheduling conflicts for a limited number of instruments or clinical procedures that do not fit within normal working hours. The problem of instrument availability is particularly acute for the C1 system. Without multiple C1 instruments, many complex experiments involving replicates, multiple time points, or multiple treatment regimens are impossible because of the difficulty of storing cells, intact, for later isolation and analysis. Treatments enabling the storage of cells in bulk for several days without degradation of RNA, while maintaining the ability to assess viability at the point of initial sample collection, would make it possible to conduct multi-sample experiments on the C1 and other platforms.

Here, we describe the adaptation and testing of a cell-permeable, reversible cross-linking fixative, dithio-bis(succinimidyl propionate) (DSP; Lomant’s reagent), as a cell preservative for single-cell transcriptomic analysis. DSP has been described previously as a reversible fixative for tissue samples preserving their integrity for immunostaining, laser microdissection, and RNA expression profiling with microarrays (3).

## Methods

### Fixation protocol

*Follow these steps to prepare and use DSP for single-cell fixation.*

1. Prepare a 50× stock solution of DSP (50mg/ml) in 100% anhydrous DMSO.
2. Dispense the stock into 100μl aliquots and store at -80°C.
3. Dilute the 50× DSP stock solution to its working concentration (1mg/ml) with PBS immediately before use, as follows:

a. In a 15ml Falcon tube, add 490μl PBS to 10μl DSP stock dropwise using a 200μl pipette while vortexing.
b. Check to ensure minimal precipitation; in case of substantial precipitation, start the dilution again with a new DSP stock aliquot.
4. Filter the 1× DSP using a 30μm filter (Miltenyi, Pre-Separation Filters; 30μm).
5. Place 1× DSP on ice.
6. Dispense 200,000 cells into a 1.5-ml Eppendorf tube.
7. Pellet cells by centrifuging for 5min at 200×g and remove supernatant.
8. Wash cells by resuspending in 200μl PBS, centrifuging, and removing supernatant.
9. Repeat PBS wash.
10. Resuspend the cell pellet gently with 200μl 1× DSP and incubate at room temperature for 30min.
11. To quench the crosslinker, add 4.1μl of 1M Tris HCl, pH 7.5 (final concentration 20mM) and mix gently by pipetting.
12. Store fixed cells at 4°C until they can be processed.

A video of the fixative preparation and DSP fixation process, showing the steps necessary to avoid precipitation of the DSP compound, is available at <u>https://youtu.be/L2aiw14IXU4</u>.

### Fluorescence staining protocol

K562 cells (ATCC^®^ CCL243^™^) were stained with one of the three staining combinations detailed below and referred to hereafter as Experiments 1, 2, and 3. In Experiment 1, all cells were stained with Hoechst 33342 “ThermoFisher” (2μM final concentration, 20 minutes at room temperature) and propidium iodide “ThermoFisher” (3.75μM final concentration, 20 minutes at room temperature). In Experiments 2 and 3, half the cells were stained with CellTracker Green CMFDA “ThermoFisher” (1μM final concentration, 30 minutes at room temperature) and LIVE/DEAD^®^ Fixable Red Dead Cell Stain “ThermoFisher” (per manufacturer protocol), while the other half were stained with CellTracker Orange CMRA “ThermoFisher” (1μM final concentration, 30 minutes at room temperature) and LIVE/DEAD^®^ Fixable Red Dead Cell Stain “ThermoFisher” (per manufacturer protocol). Post-staining, cells were thoroughly washed with PBS to remove unincorporated stain and re-suspended in PBS or 1x DSP for fresh and fixed cells respectively.

### Single-cell K562 workflow

Fresh or fixed K562 cells (ATCC^®^ CCL243^™^) were captured on the C1 system (Fluidigm) and processed using the SMARTer chemistry (SMARTer^®^ Ultra^™^ Low RNA Kit for Illumina^®^ Sequencing, Takara Clontech), according to the Fluidigm protocol, “Using C1 to Generate Single-Cell cDNA Libraries for mRNA Sequencing”, <u>PN 100-7168 Rev H1</u>. The protocol was modified for fixed cell runs to incorporate a reverse crosslinking step as described - lysis Mix A was prepared with 10.5μl Clontech Dilution Buffer plus 1μl 1M DTT rather than 11.5μl Clontech Dilution Buffer. A subset of cDNA samples was run on Agilent 2100 Bioanalyzer (High Sensitivity DNA Analysis Kit as per manufacturer’s protocol). cDNA samples were selected after analysing the cell images from the C1 integrated fluidic circuits (IFCs), and prepared for sequencing using the Nextera XT DNA Library Prep Kit (Illumina) with our own in-house primers (4). Up to 96 libraries were sequenced per experiment on a single Illumina HiSeq2500 100bp paired-end sequencing lane in Experiment 1, or Illumina HiSeq4000 75bp paired-end sequencing lane in Experiments 2 and 3.

### RNA extraction and library preparation from bulk samples

Additionally, bulk samples of 4000 K562 cells (fresh or DSP-fixed) were prepared simultaneously with the C1 run, as positive controls in each experiment. Both fresh and fixed cells were first incubated at 37°C for 30min in 50mM DTT (DL-Dithiothreitol; Sigma, D9779-5G). RNA was extracted using the RNeasy^®^ Plus Micro Kit (Qiagen) and eluted in 14μl of RNAse-free water. A total of 1μl aliquots of the extracted RNA were processed using portions of the same RT and PCR reagent master mixes described for the tube controls in the C1 protocol above, followed by Nextera XT Library Generation. Nextera XT libraries from bulks were pooled and sequenced together with the single-cell libraries.

### RNA sequencing, mapping, gene counts and QC

RNA-seq reads were trimmed for Nextera and Illumina adapter sequences using skewer-v0.1.125 (5). Trimmed reads were mapped to a modified reference genome comprising the human genome *Homo sapiens* GRCh37 (human_g1k_v37, available at <u>http://software.broadinstitute.org/software/genomestrip/node_ReferenceMetadata.html;</u> last accessed 20 August 2017) and ERCC RNA Spike-In Mix sequences (ThermoFisher), using HISAT2 version-2.0.0-beta(6) with default parameters. Duplicate reads were marked using MarkDuplicates.jar implemented in Picard tools v1.92 (<u>http://broadinstitute.github.io/picard/</u>). BAM alignments were name sorted with Samtools version 1.1(7). Alignment metrics were calculated using CollectRnaSeqMetrics from Picard tools for full BAM files and with potential PCR duplicates marked. Reads mapping uniquely to genes annotated in ENSEMBL release 75 (8) were counted using featureCounts (9) implemented in subread-v1.5 (10). The distribution of reads among several categories – assigned reads (mapped uniquely to exons), multiple mapping, ambiguous mapping, no feature (mapped uniquely to intronic and intergenic regions) – was obtained from featureCounts summary. All further metrics were calculated using R core tools, version 3.2.4 (11). Read counts were normalized to counts per million (CPM) and the numbers of detected genes per sample were calculated by counting genes with at least 1 CPM. Unsupervised clustering of single cells based on gene expression values was performed using the consensus-clustering algorithm implemented in Bioconductor package, SC3 version 1.1.4 (12).

## Results and Discussion

### Cell imaging

The DSP fixation protocol was initially optimized for cell appearance, RNA integrity, and cDNA profile, using bulk cell samples. Using our protocol above, DSP-fixed cells showed similar morphology to unfixed cells (Figure 1), with no obvious clumping or shrinkage upon treatment. In case of cells stained using common dyes such as Hoechst 33342 or propidium iodide (PI), the cell-stain was retained at least up to 8 days in both fresh and fixed cells (Figure 2).

**Figure 1.**
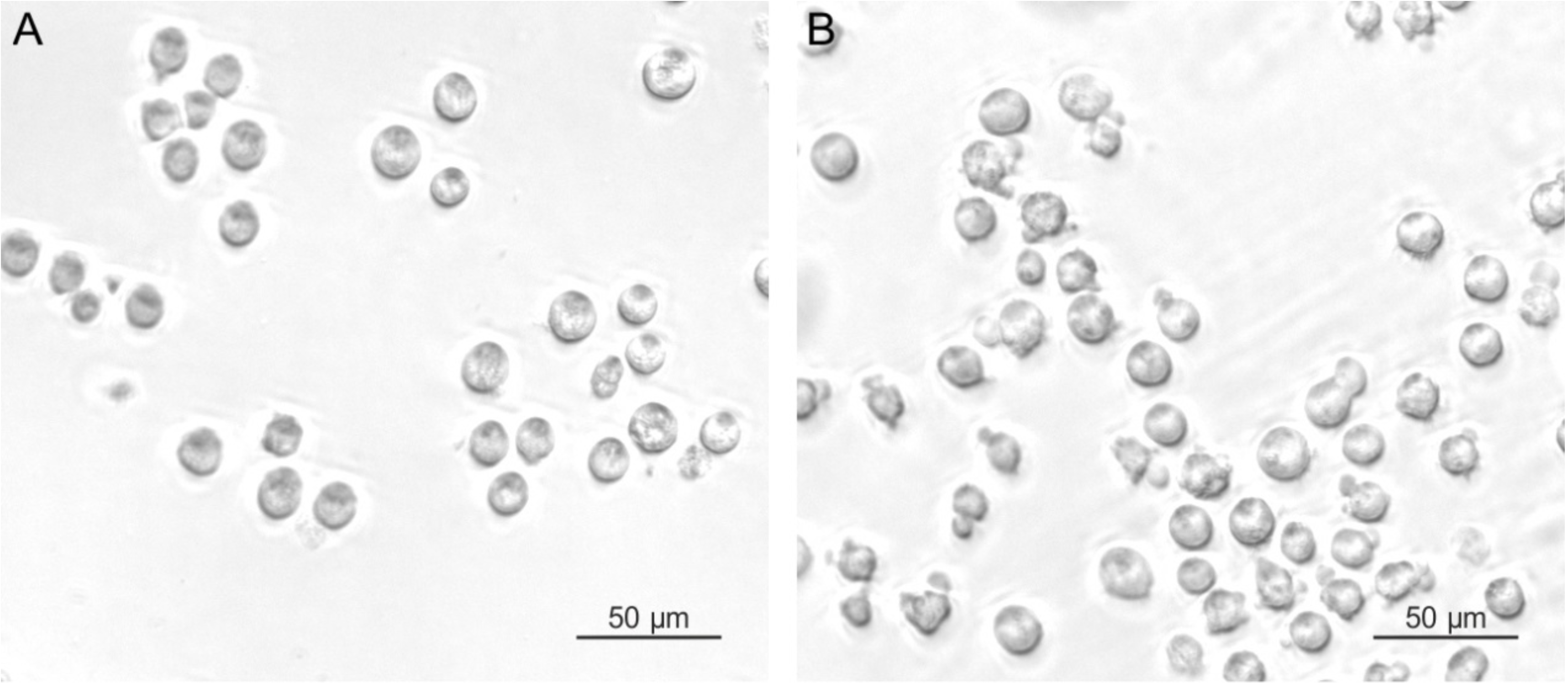
Morphology of K562 cells before (A) and after (B) DSP treatment. DSP fixation did not cause detectable shrinkage or clumping.

**Figure 2.**
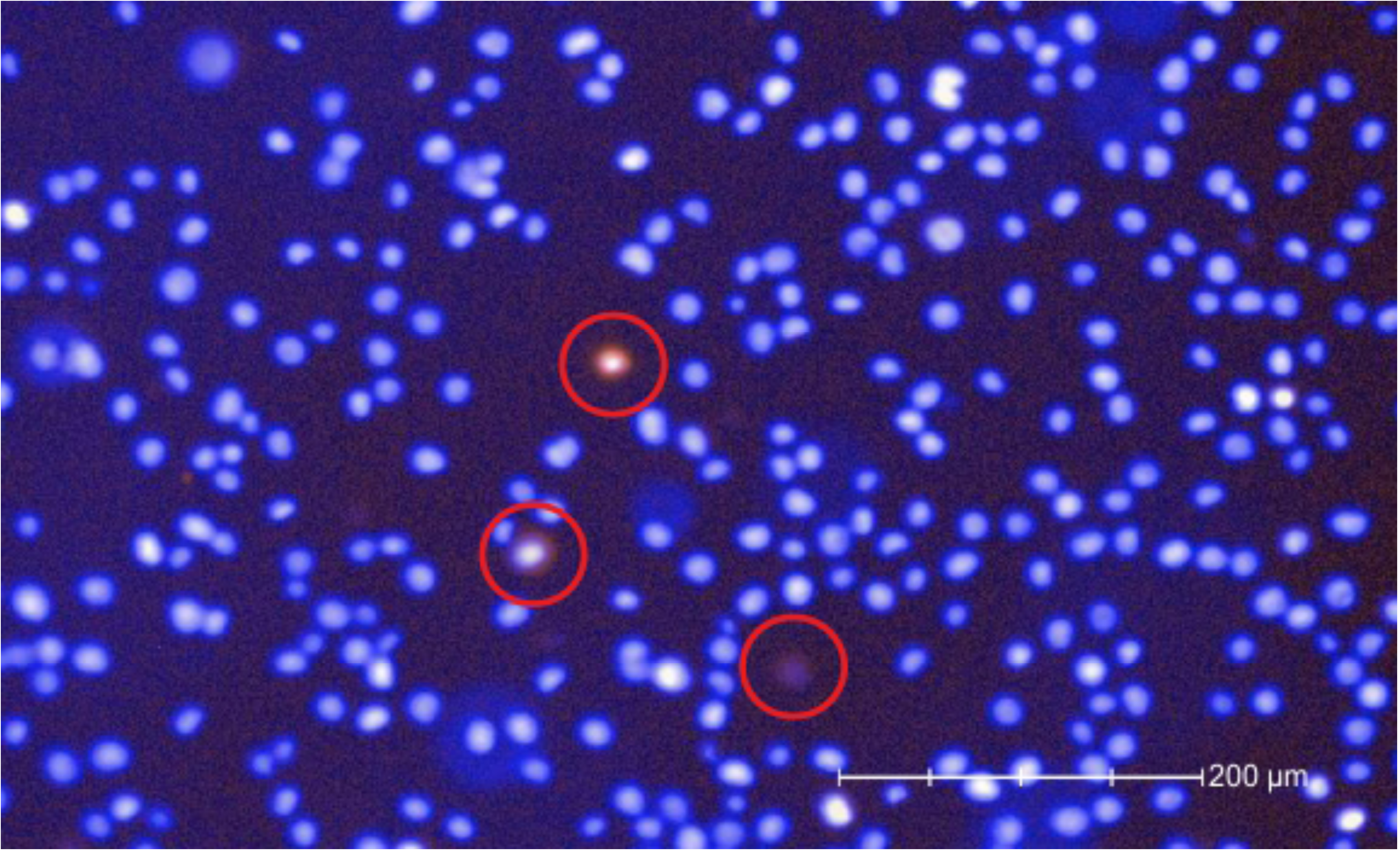
Fluorescent staining of K562 cells after DSP treatment. In an overlay image of fixed K562 cells stained with Hoechst 33342 (blue) and propidium iodide (PI; red) before DSP treatment and then photographed 1 hour after fixation. Double-stained cells are circled in red.

### Cell capture

K562 cells were captured on the C1 system in three separate experiments that compared fresh cells - captured immediately, without DSP treatment - with cells captured 3 and 7 days (Experiment 1) or 2 days (Experiments 2 and 3) after fixative treatment. This feature of the method in principle enables the collection of stain-related data by imaging fixed cells in the C1 IFC in the same manner as for fresh captured cells, including resolution of nuclei in capture sites, tracking of differentially stained sub-populations in experimentally mixed samples, and “freezing” viability status at the time of fixation. Single fixed cells were readily captured in C1 IFCs with no apparent difference in capture rate compared with fresh cells (Figure 3).

**Figure 3.**
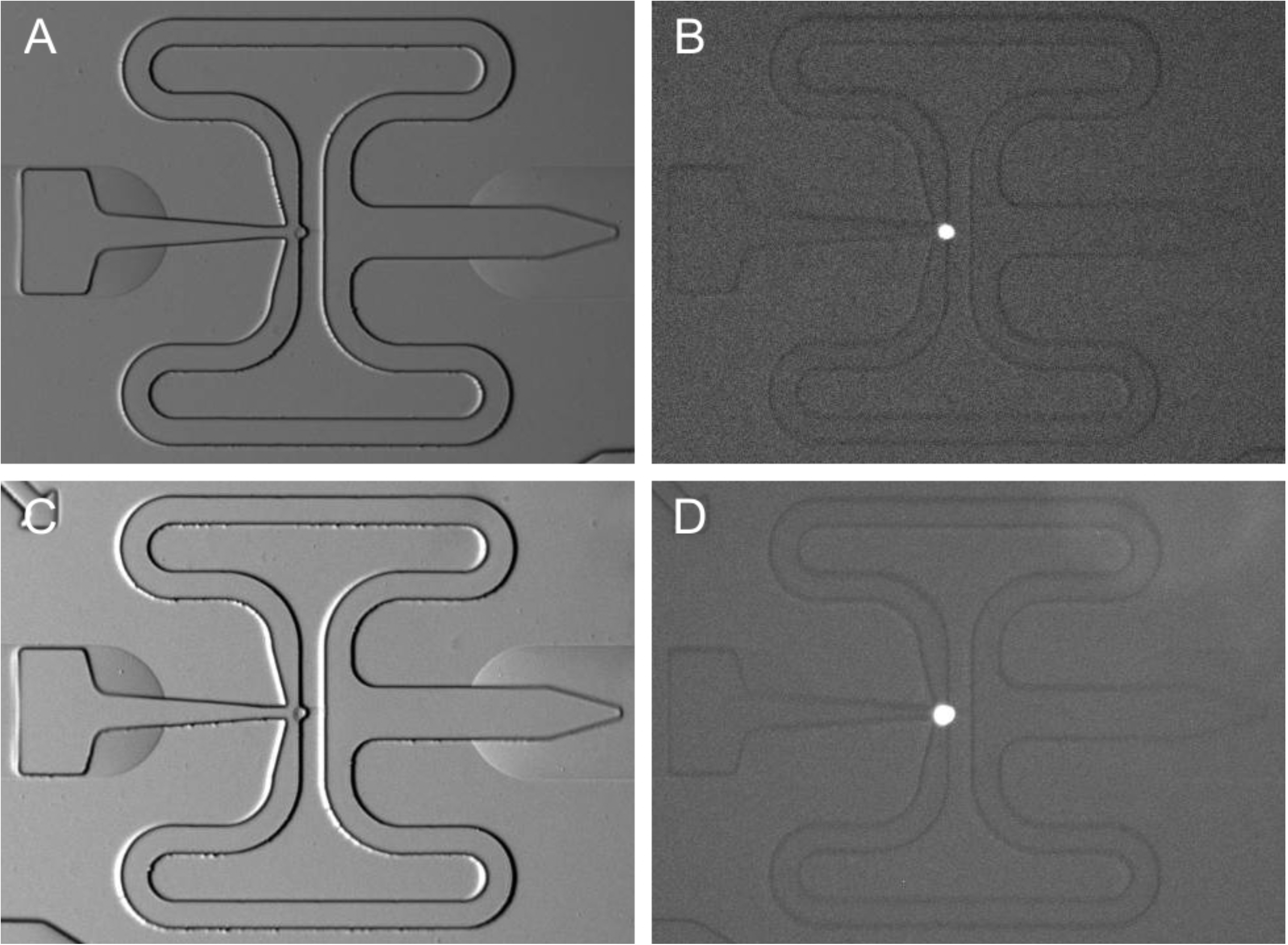
Imaging of captured DSP-fixed K562 cells. Cells were captured on a Fluidigm C1 microfluidics IFC 3 days (A and B) or 7 days (C and D) after fixation, and imaged under bright field (A and C), or fluorescence for Hoechst 33342 (B and D).

### cDNA yield

The cDNA profile did not differ significantly between single cells captured fresh and at different times after fixative treatment (C1 single-cell data from Experiment 1, Figure 4). There was noticeably higher variation in cDNA yield among fresh than among fixed cells (Figure 5). As expected, non-viable (PI+) cells tended to have low yield and, intriguingly, there appeared to be very few such cells captured from fixed K562 samples. This finding suggests a potentially important benefit of fixation, namely, by limiting the handling of live cells, treatment with DSP before the capture step may better preserve the original biological state of analysed cells, compared to fresh cells.

**Figure 4.**
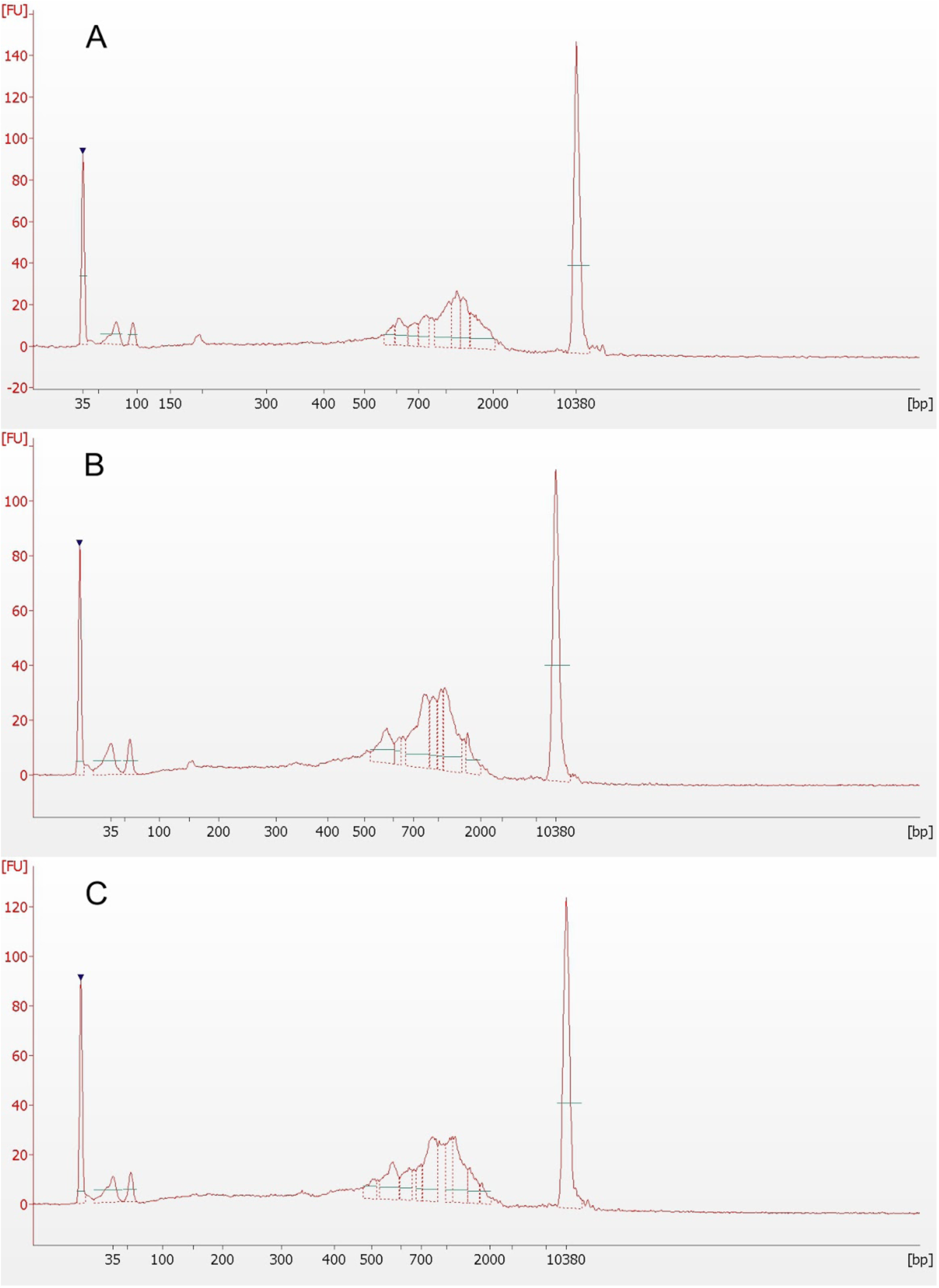
cDNA profiles of individual C1-captured K562 cells. Bioanalyzer (Agilent) profiles of cDNA from typical high-yielding cells revealed little or no effect of fixation. Cells were isolated: (A) fresh, without fixation; (B) 3 days after DSP treatment; and (C) 7 days after DSP treatment.

**Figure 5.**
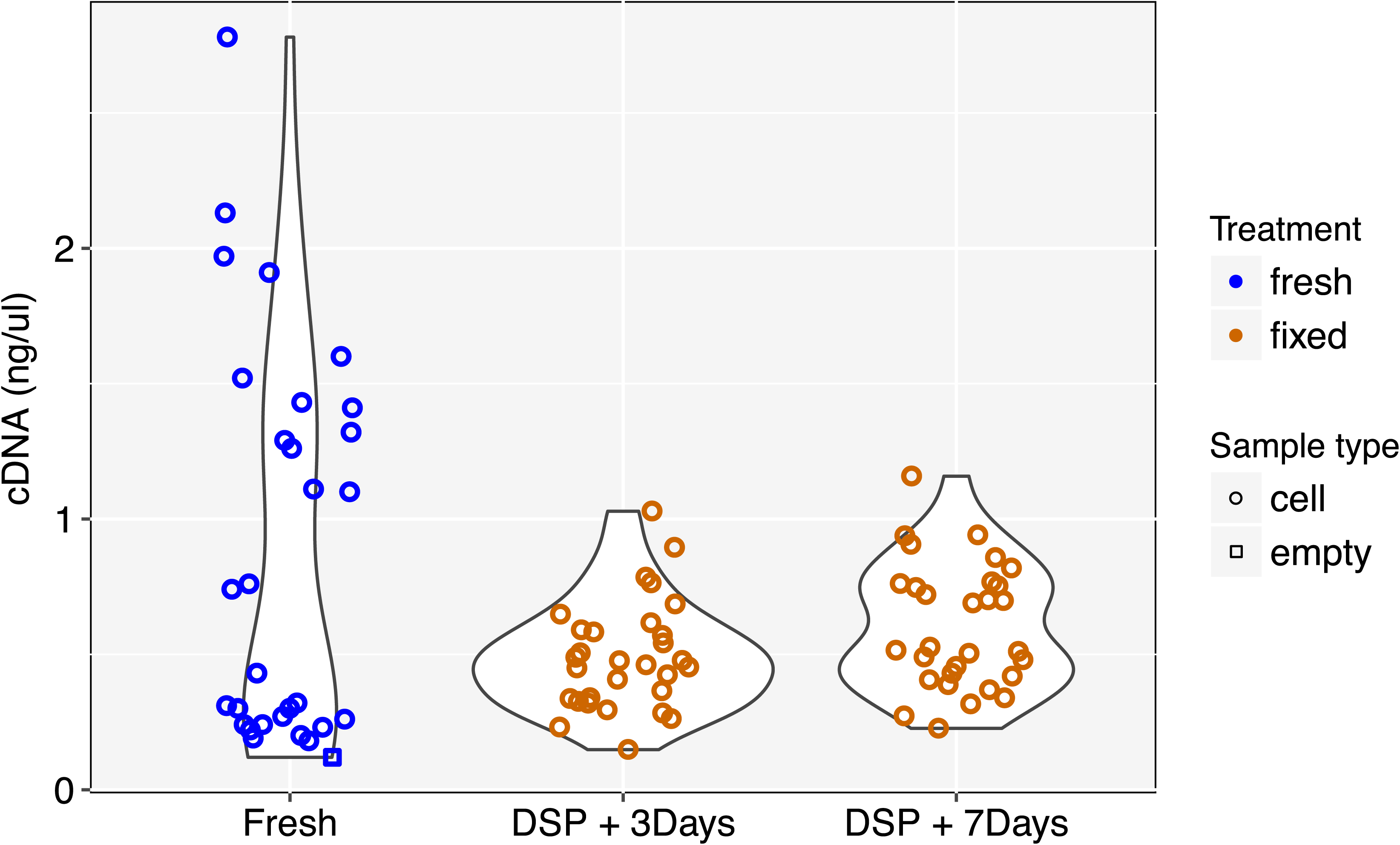
cDNA yields of individual C1-captured K562 cells. Cells were isolated fresh (without fixation) or 3 or 7 days after DSP treatment. Mean yield differed between C1 runs, but appeared to be more variable in fresh samples.

### Bulk sequencing comparison of fresh and fixed K562 cells

We included RNA prepared from bulk cells as positive controls and empty wells as negative controls. We see high pairwise correlation for overall gene expression (Pearson’s R > 0.93, p < 0.01) between fresh and fixed bulk samples (Experiment 1: Figure 6). The correlation between fixed samples from the same experiment stored for 3 days and those stored for 7 days after fixation is comparable to that between fresh bulk samples (R ∼ 0.97, p < 0.01).

**Figure 6.**
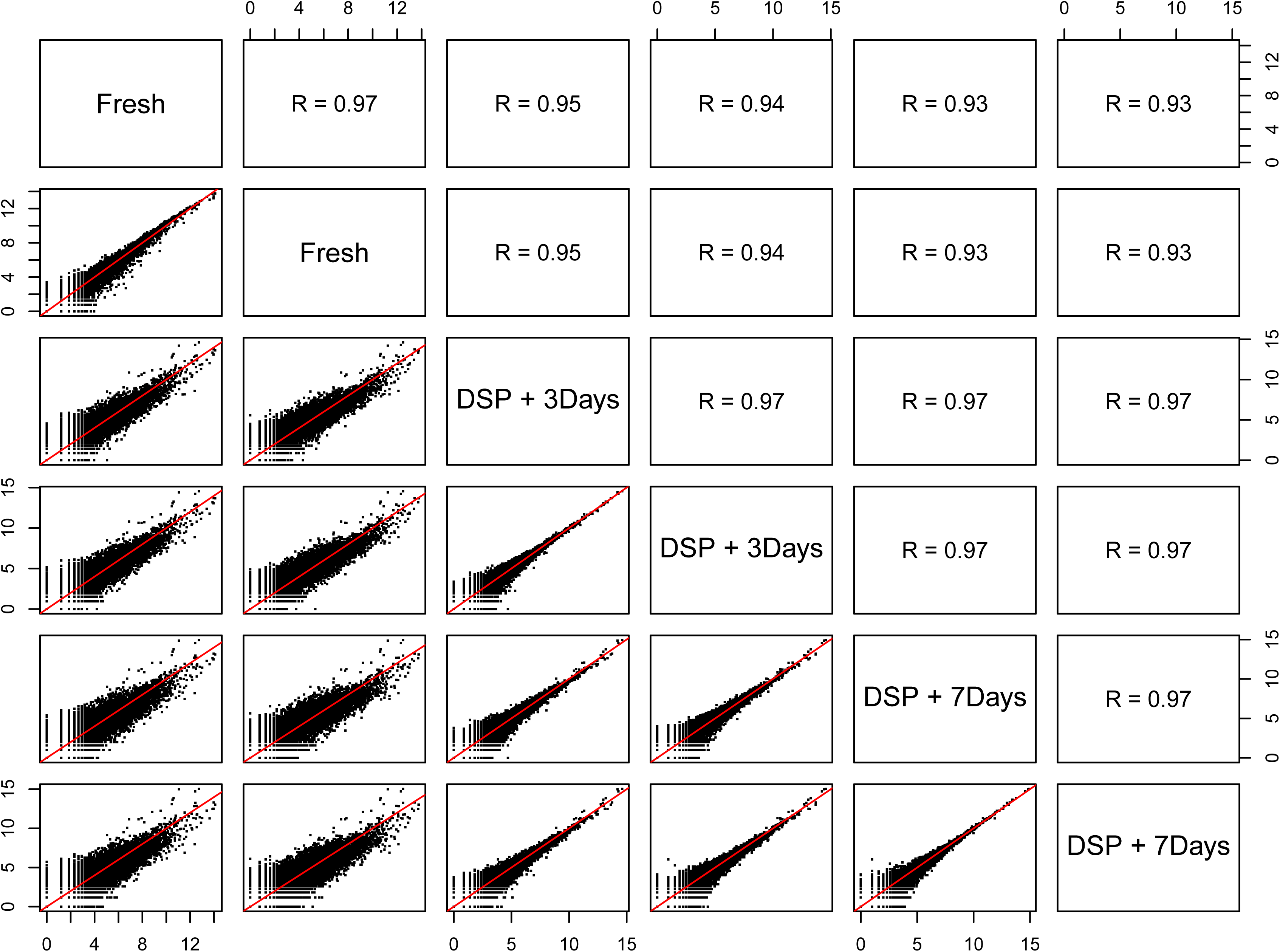
Pairwise correlation in bulk gene expression. Fresh and fixed K562 cells had highly similar patterns of gene expression (above the diagonal: Spearman’s rank correlation coefficient; below the diagonal: scatterplots of log_2_ transformed gene counts). The relative similarity of fixed cells stored for different times reveals signs of a systematic effect of fixation.

### Library complexity is maintained for fixed single cells

With approximately 5000-7000 genes detected and approximately 40-60% reads mapping to the most abundantly expressed 500 genes in the cell, both fresh and fixed K562 cells from Experiments 1-3 produced libraries of the expected complexity (Figure 7). A subset of low-quality libraries with relatively few distinct transcripts and low cDNA yields (lower right of plots in Figure 7), characteristic of PI+ dead cells and empty capture sites, was evident among fresh cells, but notably not among fixed cells. These data were removed from subsequent analyses. The numbers of detected coding genes were not distinguishable between the remaining fresh and fixed cells (p > 0.1, Mann-Whitney U test).

**Figure 7.**
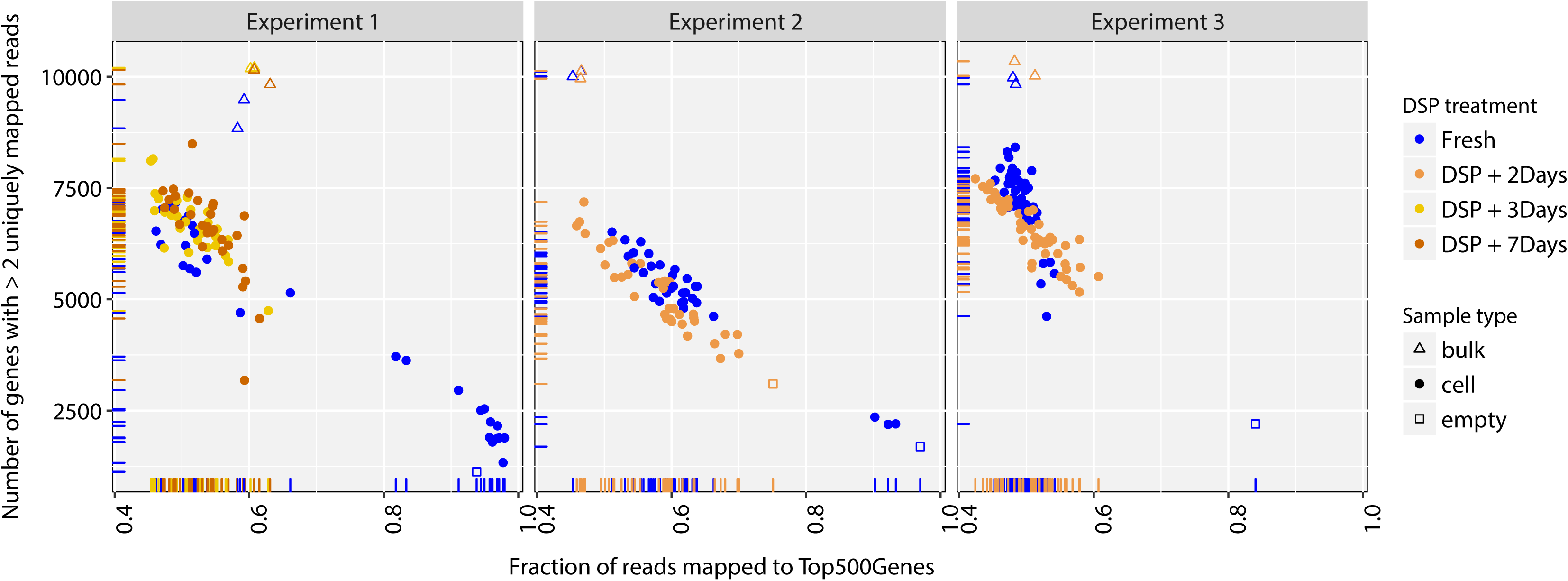
Transcript complexity: Counts of detected genes vs fraction of reads in top 500 genes. Measures of the recoverable complexity of transcripts show little difference between fresh and fixed cells, except that there were more propidium iodide+ fresh cells in each experiment, which produced lower cDNA yields and, if sequenced, failed on standard complexity metrics.

### Gene body coverage

We used SMARTer chemistry, designed to retrieve full-length transcripts as cDNA, in this study. Plotting the relative coverage of reads along the (normalised) length of all detected genes for each cell (Figure 8A) reveals the expected lack of 5’-to-3’ bias in high-quality fresh cells. In fixed cells, however, there is a mild bias in coverage towards the 3’ ends of genes, which appears to increase with time of storage after DSP treatment. The bias is also present in bulk samples (Figure 8B) and is different in appearance from the irregular (‘spiky’), 3’-biased coverage characteristic of degraded RNA that we have observed in low-quality cells or empty wells (Figure 8C). The absence of a reduction in gene-level complexity that accompanies the observed bias suggests that, while DSP treatment and storage may affect the retrieval of whole transcripts from cells, the priming efficiency of reverse transcription may be comparable to that in fresh cells and the method may be particularly suited to end-counting RNA-Seq methods.

**Figure 8.**
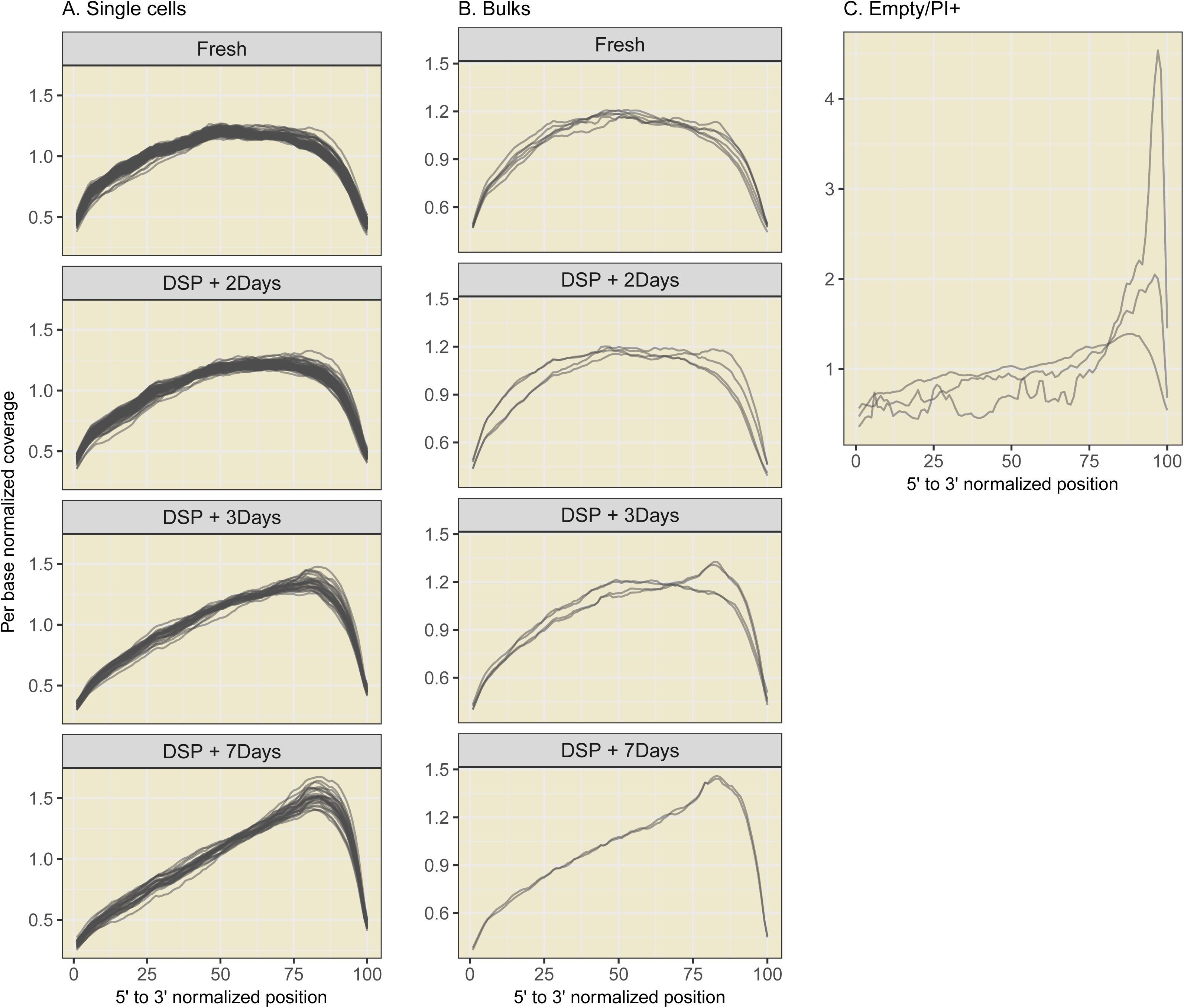
Biased sequencing coverage across the gene body. Relative read-depth across all genes, normalised to the same length, reveals consistently greater bias in fixed cells than in fresh cells (A. Single cells, B. Bulks), which appeared to increase with time of storage after DSP treatment. Gene-body coverage profile for cells with low quality/complexity or empty wells show spiky coverage with very high 3’ bias (C.) Only cells passing quality/complexity filters were included.

### Experimental heterogeneity in cell-type profiles

We compared single-cell expression profiles across several experiments and C1 runs to illustrate and contrast heterogeneity at several levels: within a sample of cells, between methods and storage time-points for the same sample, and between experiments. In comparisons between all K562 cells from three experiments (Figure 9), the dominant factor distinguishing cells appears to be experimental batch (note: a distinct lot of K562s was used in Experiment 1 vs. Experiments 2 and 3), rather than whether the cells were fresh or fixed.

**Figure 9.**
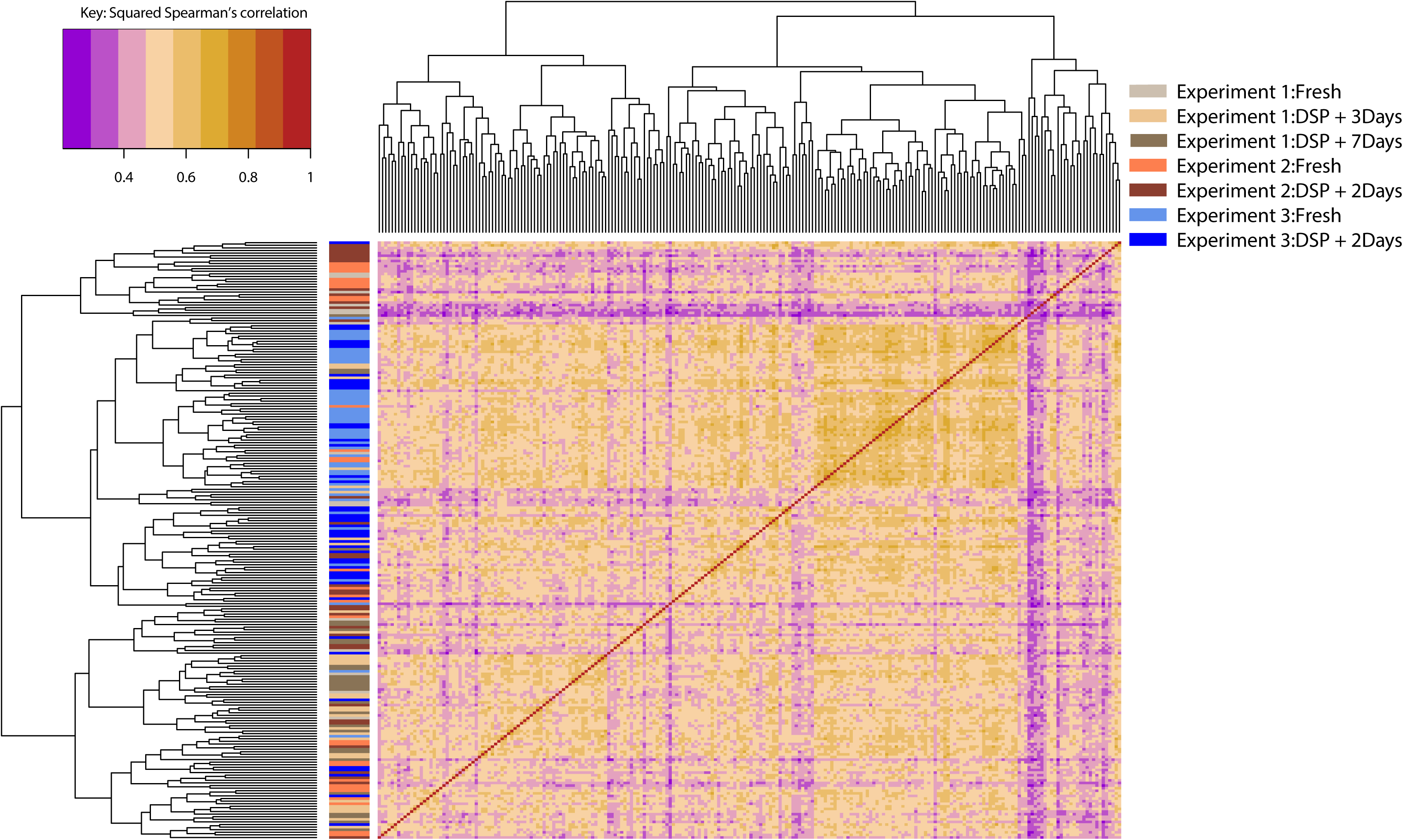
Pairwise expression correlation for all K562 cells. Heatmap of squared Spearman’s correlation coefficient of the expression of the top 500 genes observed across all experiments shows high similarity of expression profiles between fresh and fixed cells, especially for Experiment 3.

Unsurprisingly, K562 cells can be easily and reliably distinguished from primary cells of a very different type, intestinal stromal cells from colonic biopsy samples (Figure 10). The observation in these clustering comparisons that fixed cells are often interspersed amongst fresh cells from the same experiment implies that the technical effects of DSP treatment on the observed transcriptome, at the gene level, may be of a smaller magnitude as batch effects. This is good news for a researcher seeking to replace multiple experiments on different days with a single experiment in which fixed cells are processed in successive runs across several days.

**Figure 10.**
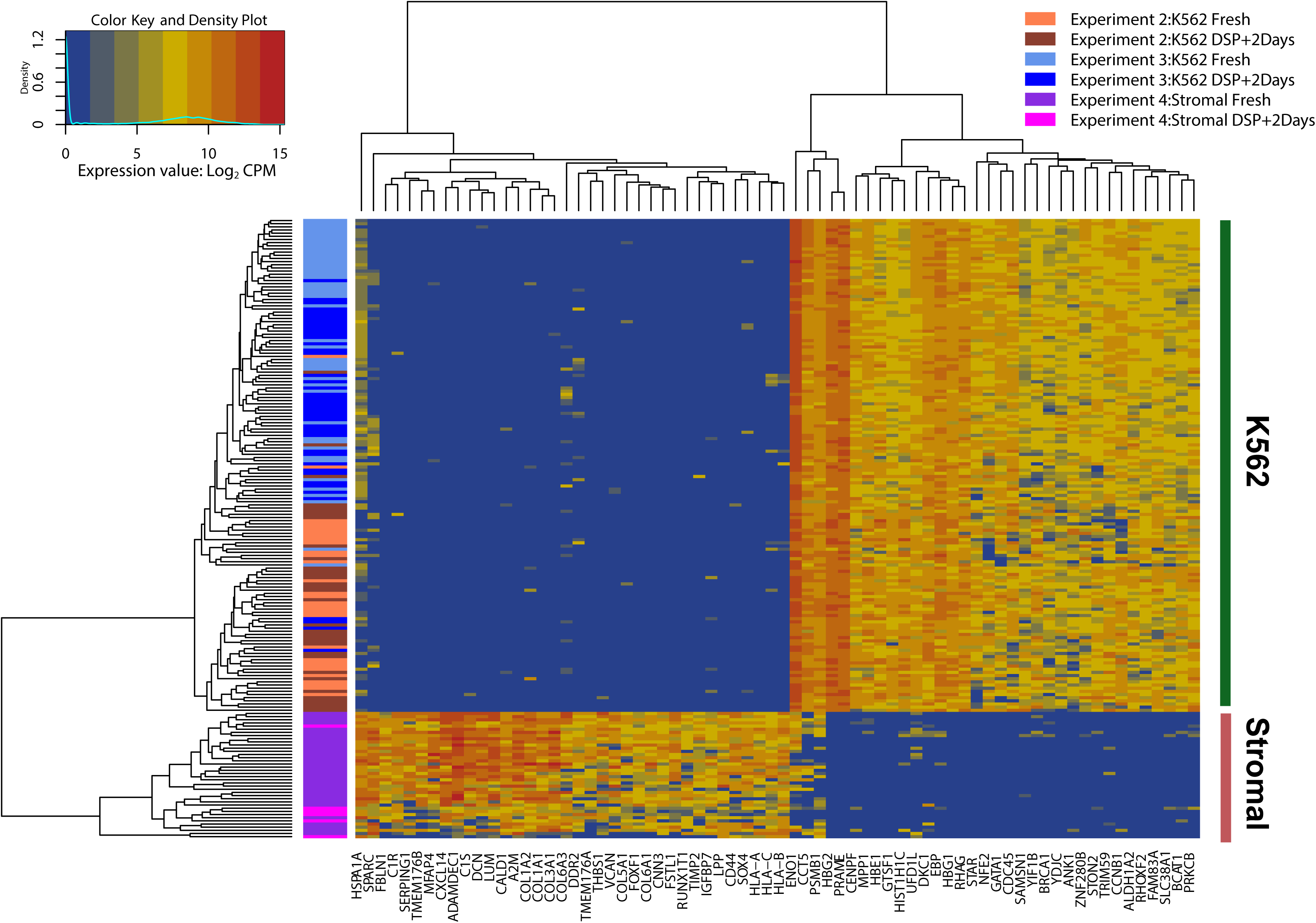
Unsupervised clustering comparison of human intestinal stromal cells and K562 cells. Along the y-axis, cells are arranged by highest similarity, revealing two clear clusters, and cells of the same type with different treatments are interspersed. The x-axis shows the top 35 marker genes for each cluster, corresponding to a cell type, identified by SC3 clustering.

## Conclusions

Single-cell genomics is leading to tremendous advancements in our understanding of the complexity of cellular models and cellular systems. Such studies may require the simultaneous production of many samples, whose single-cell analysis can be constrained by distance to and throughput of specialist facilities when cells need to be isolated and processed immediately to avoid sample degradation.

We have demonstrated that DSP treatment can be used for preserving cells for subsequent cell isolation and processing for single-cell RNA sequencing for at least several days after sample collection. Preserved cells are amenable to microfluidic manipulation and can be stained with most commonly used dyes for tracking and viability assessment. RNA-seq of fixed cells, captured and processed up to 7 days after DSP treatment, shows highly similar transcript complexity and transcriptome identity to fresh cells.

Cells preserved with DSP are stabilized at the time of sample collection and large numbers of fixed samples can be easily transported and subsequently processed for RNA-seq. This separation of the place and time of sampling from downstream processing enables complex study designs for single-cell experiments. DSP preservation has traditionally been applied to cell solutions as well as tissues, and DSP-treated samples are amenable for immuno-staining and FACS sorting(13, 14). Therefore, our method for cell preservation may also have utility for preservation of tissues at sampling before dissociation, and in combination with FACS sorting of stabilized cells.

## Data availability

The RNAseq data discussed in this publication have been deposited in NCBI’s Gene Expression Omnibus (15). The sequencing data, experimental metadata and QC metrics are accessible through GEO Series accession number GSE98734 (http://www.ncbi.nlm.nih.gov/geo/query/acc.cgi?acc=GSE98734).

## Consent

Colonic biopsy samples were collected, following written informed consent, from patients attending for clinically indicated endoscopy procedures at Oxford University Hospitals NHS Foundation Trust. Ethical approval for the study was obtained from NRES Committee Yorkshire & The Humber - Leeds West (REC reference: 14/YH/1116).

## Author contributions

KJL and RB conceived and oversaw the study. KJL, SL and MA developed the methodology. RB, MA and ES designed the experiments. MA, CB, LC and JK undertook the experiments. ES, MA, HL and RB analysed the data. RB, ES, MA and KJL prepared the manuscript. PP, AS and DB contributed to the conception of the study and the experimental design. All authors were involved in the revision of the draft manuscript and have agreed to the final content.

